# Improved *Apis mellifera* reference genome based on the alternative long-read-based assemblies

**DOI:** 10.1101/2021.04.30.442202

**Authors:** Milyausha Kaskinova, Bayazit Yunusbayev, Radick Altinbaev, Rika Raffiudin, Madeline H. Carpenter, Alexey Nikolenko, Brock A. Harpur, Ural Yunusbaev

## Abstract

*Apis mellifera* L., the western honey bee is a major crop pollinator that plays a key role in beekeeping and serves as an important model organism in social behavior studies. Recent efforts have improved on the quality of the honey bee reference genome and developed a chromosome-level assembly of sixteen chromosomes, two of which are gapless. However, the rest suffer from 51 gaps, 160 unplaced/unlocalized scaffolds, and the lack of 2 distal telomeres. The gaps are located at the hard-to-assemble extended highly repetitive chromosomal regions that may contain functional genomic elements. Here, we use *de-novo* re-assemblies from the most recent reference genome Amel_HAv_3.1 raw reads and other long-read-based assemblies (INRA_AMelMel_1.0, ASM1384120v1, and ASM1384124v1) of the honey bee genome to resolve 13 gaps, five unplaced/unlocalized scaffolds and, the lacking telomeres of the Amel_HAv_3.1. The total length of the resolved gaps is 848,747 bp. The accuracy of the corrected assembly was validated by mapping PacBio reads and performing gene annotation assessment. Comparative analysis suggests that the PacBio-reads-based assemblies of the honey bee genomes failed in the same highly repetitive extended regions of the chromosomes, especially on chromosome 10. To fully resolve these extended repetitive regions, further work using ultra-long Nanopore sequencing would be needed. Our updated assembly facilitates more accurate reference-guided scaffolding and marker/sequence mapping in honey bee genomics studies.

## INTRODUCTION

An accurate reference genome is an important starting point in translating an organism's genomic information to its function at the molecular, cellular, and organismal levels. The genome of the western honey bee (*Apis mellifera* L., henceforth honey bee) has been a boon to our understanding of genomics in insect and eusocial species (Honeybee Genome Sequencing Consortium 2006; Harpur *et al.* 2019). The original reference genome (Honeybee Genome Sequencing Consortium 2006) was recently updated (Wallberg *et al.* 2019), providing to the community a chromosome-level assembly that is more contiguous and complete than the previous reference assembly (Elsik *et al.* 2014). Unfortunately, it still has a number of issues that hinder downstream genomic inferences. Specifically, the new reference has 51 unsolved genomic gaps, 2 lacking distal telomeres (Figure 1), and 160 unplaced/unlocalized scaffolds. There are 17 arbitrary gaps of 25 and 200 bp in the Amel_HAv_3.1, and the remaining varies from 393 to 345,148 bp. There are 14 gaps located within the genes of the Amel_HAv_3.1. The distal telomeres of the Amel_HAv_3.1 are assembled, except for chromosomes 5 and 11. In addition to these gaps, there are several problematically assembled regions in chromosomes 3, 6, 7, 10, and 11, which demonstrate significantly higher levels of reads coverage variation (Figure 1).

**Figure 1.**
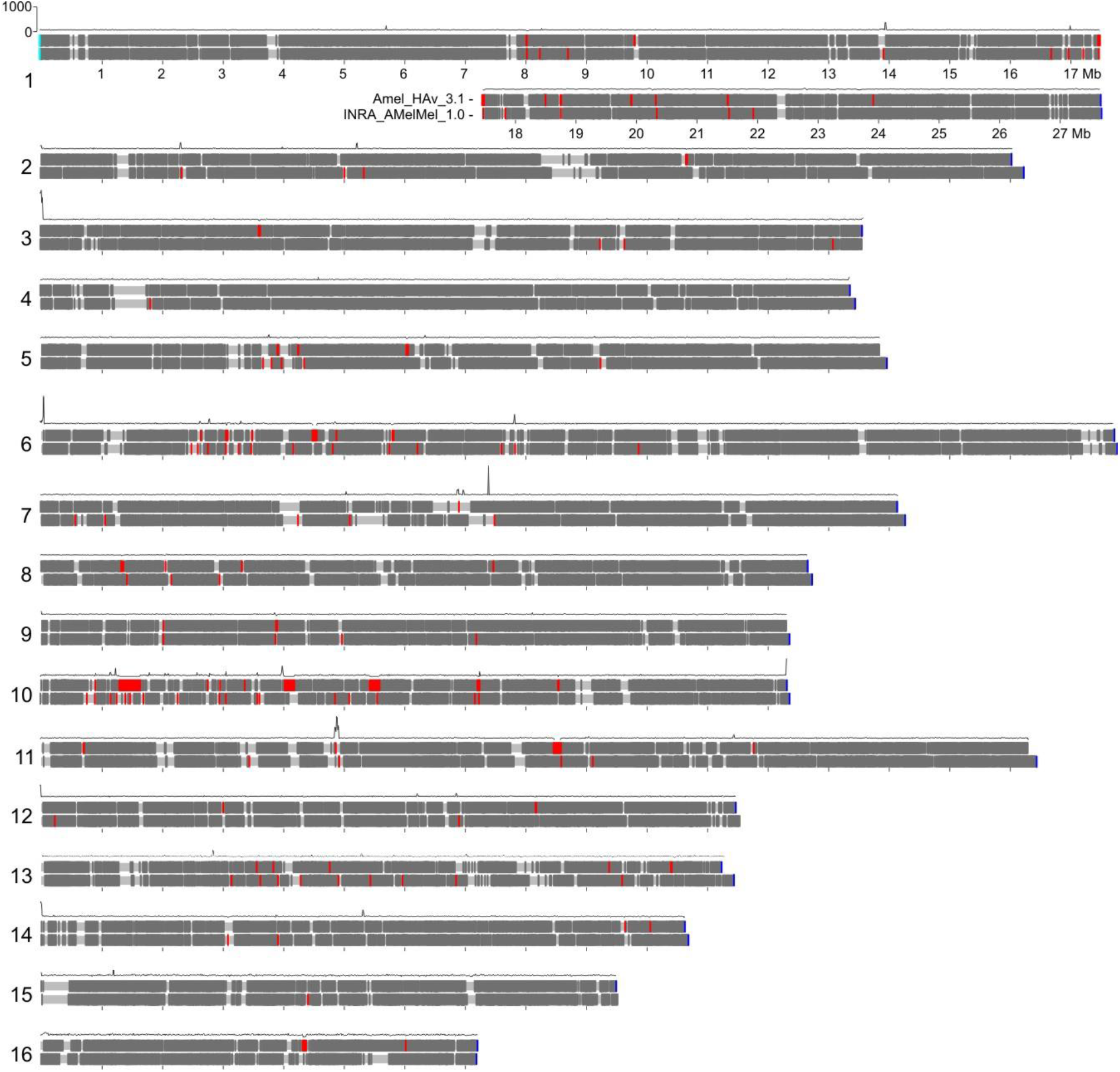
Ideograms of two assemblies of the *A. mellifera* genome Amel_HAv_3.1 (upper) and Amel_INRA_1.0 (lower) with the mapped genes (dark gray), telomeric TTAGG (blue) and CCTAA (cyan) motifs, polyN gaps (red), and Amel_HAv_3.1’s PacBio reads coverage (black curve).

Identifying the sequences that fill the genomic gaps could facilitate the discovery of novel genomic features in the honey bee genome that can lead to important biological insights and would improve downstream genomic analysis. For example, closed gaps in the human reference genome were found to be enriched in repetitive elements and contain functional genomic elements (Zhao *et al.* 2020). There has been considerable progress in developing gap closing methods in the past decade, such as methods based on the local assembly approach (English *et al.* 2012; Bayega *et al.* 2020; Miga *et al.* 2020) and the assembly-to-assembly approach (Thomma *et al.* 2016; Shi *et al.* 2016; Zhao *et al.* 2020). These methodological advancements allowed significant progress in resolving gaps in the human reference genome. Unlike the progress with the human genome, there are still issues regarding the gaps in the honey bee reference.

Here, we sought to improve the current assembly by filling in the remaining gaps and developing a telomere-to-telomere chromosomal reference sequence. We use two *de-novo* re-assemblies from Amel_HAv_3.1 PacBio reads, referred to as “re-assemblies”, and three *de-novo* assemblies from PacBio reads derived from different honey bee subspecies, referred to as “alternative assemblies”, to improve the honey bee reference genome Amel_HAv_3.1.

## MATERIALS AND METHODS

Our method (Figure 2A) utilizes five genomic datasets including the current version of the honey bee reference (Amel_HAv_3.1), two *de novo* re-assemblies of the reference, and three non-reference alternative *de novo* genome assemblies derived from the different *A. mellifera* subspecies (see “Genomic data” section below). First, we identified the coordinates of the gaps and the genes flanking them in the Amel_HAv_3.1 reference genome. Then, we determined the flanking genes' positions in alternative assemblies. The flanking genes were used as markers to find and extract the Gap Closing Sequences (GCS) from the alternative assemblies (Figure 2B). Next, we selected candidate GCSs that demonstrate the best alignment to the corresponding gap region. In addition, for each filled gap, we verified whether the PacBio raw reads from the Amel_HAv_3.1 are properly aligned to the region. If they were not, we discarded the tested GCS. All the gaps filled in our study were carefully curated manually. We also positioned unplaced scaffolds and restored lacking telomeres by comparing gene coordinates in different assemblies. All the redundant sequences were removed from the corrected assembly. Finally, we evaluated and validated the corrected_Amel_HAv_3.1.

**Figure 2.**
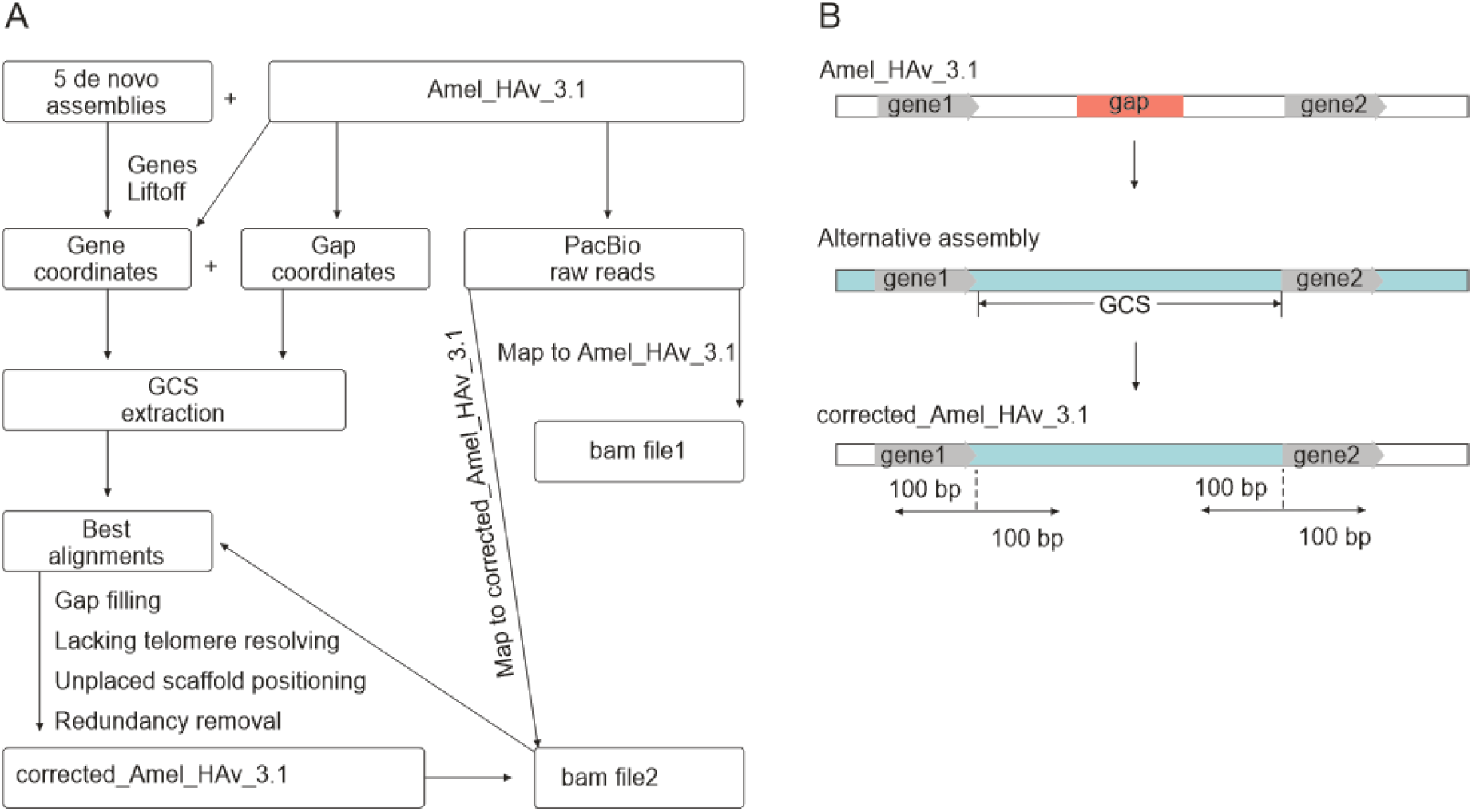
Workflow of our approach to identify and validate Gap Closing Sequences (GCSs) (A), (B) Schematic of our gap-closing approach that was used to improve the *A. mellifera* reference genome Amel_HAv_3.1.

### Genomic data

The Amel_HAv_3.1 ****reference genome**** (Wallberg *et al.* 2019) along with raw reads were downloaded from NCBI (Table S1).

****Reference de novo re-assemblies**** were built out of Amel_HAv_3.1 raw reads using two assemblers: Flye v2.8 (Kolmogorov *et al.* 2019) and NextDenovo v2.3.1 (https://github.com/Nextomics/NextDenovo). Default parameters were used except where stated. All the commands and parameters used for each tool are given in Table S2. The re-assembled contigs were ordered and oriented in RaGOO (Alonge *et al.* 2019) using Amel_HAv_3.1 as a reference. The assemblies were polished in NextPolish (Hu *et al.* 2020) using PacBio and Illumina reads. The re-assemblies from the Flye and NextDenovo are referred to as Amel_HAv3_1_reFlye and Amel_HAv3_1_reND, respectively.

***Non-reference de novo alternative assemblies*** based on SMRT PacBio long reads for *A. m. mellifera* (Assembly: INRA_AMelMel_1.0; NCBI Bioproject: PRJNA450801), *A. m. carnica* (ASM1384124v1, PRJNA644991), and *A. m. caucasica* (ASM1384120v1, PRJNA645012) were downloaded from NCBI. All assemblies based on PacBio reads were required to have coverage higher than 100.0x. To achieve chromosome-scale assembly, the ASM1384120v1 contigs were re-scaffolded using RaGOO and Amel_HAv_3.1 as a reference. The INRA_AMelMel_1.0 and ASM1384124v1 chromosome-scale assemblies were used as is.

### Gap-closing

We used genes that flank reference gaps as markers to find and extract GCSs from the alternative assemblies (Figure 2). For this, we mapped genes from the Amel_HAv_3.1 reference assembly to the alternative assemblies. Ordering and orientation of the genes were compared between these alternative assemblies and Amel_HAv_3.1 (Table S3.1-3.6). Next, we found GCSs in the queried alternative assemblies. Then, we generated three files using BEDTOOLS: (1) a fasta file of the reference genome Amel_HAv_3.1 with deleted gap regions. Gap regions were deleted from the genome based on the end (or start) position of the terminal gene, flanking the gap upstream, and start (or end) position of the first gene, flanking the gap downstream; (2) a fasta file with the GCSs from the gap-closing assembly. GCSs were also retrieved from assemblies based on the positions of the gap-flanking genes. If the gap in the reference genome was located within the gene, we pasted this gene from another assembly that contained the complete sequence of the gene; (3) a fasta file with the genomic region flanking the start and end positions of the GCS. We extracted 100 bp fragments located upstream and downstream of each GCS and aligned them to the reference assembly to check that ends of the GCSs correspond to sequences in the reference genome. The fasta files (1) and (2) were merged and GCSs were pasted in corresponding regions manually. To validate GCSs, we aligned the 200 bp fragments located upstream and downstream of each GCS to the reference assembly (Figure 2B). Then, we mapped Pacbio raw reads to the corrected Amel_HAv_3.1 and calculated genome coverage. The same approach was used to recover telomeres in chromosomes 5 and 11.

### Gene annotation liftoff

We used the Liftoff software (Shumate and Salzberg 2020) to map the genes from the Amel_HAv_3.1 reference to the re-assembled and alternative assemblies.

### Assembly assessment

Assembly statistics were computed using Quast (Table S4). We used BUSCO v. 4.1.2 (Waterhouse *et al.* 2019) and Liftoff to assess gene sets in honey bee assemblies. Minimap2 (Li 2018) was used to map Pacbio reads to the initial and corrected Amel_HAv_3.1 assembly (minimap2 -ax map-pb). To calculate genome coverage, we used CLC Genomics Workbench 20.0 (https://digitalinsights.qiagen.com) and Samtools (samtools depth -a, https://www.htslib.org/).

### Computing resources

All the programs were run on the WorkStation HP Z-series and Dell PowerEdge T-series with 6 core processors and 196Gb RAM in total. Also, we used the public server at usegalaxy.org (Sloggett *et al.* 2013) to run BUSCO and Quast.

#### Data availability

The assembly generated in this study and supplementary materials are available at the Figshare repository from https://figshare.com/s/c8d6c0291893405d4409.

## RESULTS AND DISCUSSION

### Gap-closing in the Amel_HAv_3.1 reference genome

We selected 11 GCSs from the two Amel_HAv_3.1 re-assemblies and three long-read alternative assemblies. In case of choice between the re-assembled Amel_HAv_3.1 and alternative assemblies, we preferred the first one. And in case of choice between alternative assemblies, we selected the one that gave the best genome coverage with PacBio reads.

Altogether, we closed 9 gaps in the Amel_HAv_3.1 reference using our re-assembly approach: gaps 4, 6, 8, and 9 in chromosome 1; gap 1 in chromosome 2; gaps 3 and 4 in chromosome 8; gaps 1 and 2 in chromosome 16. Five of these closed gaps were located within genes and three gaps were in intergenic regions. We also found that the gap 4 in chromosome 1 arose due to low sequencing coverage in the region.

We found that the gap-containing regions that we processed using our GCSs were enriched for repeats (Figure S1). These repetitive elements probably hindered previous assemblies and resulted in gaps. In these regions, we observed discrepancies in the ordering and orientation of the genes for different assemblies (Figures S2.1 and S2.1a). The details of the remaining gaps that we corrected in this study are given in Supplementary (Figures S2.2-2.10). In Figure 3, we show the corrected exon-intron structure of the LOC410785 gene before and after the gap closing.

**Figure 3.**
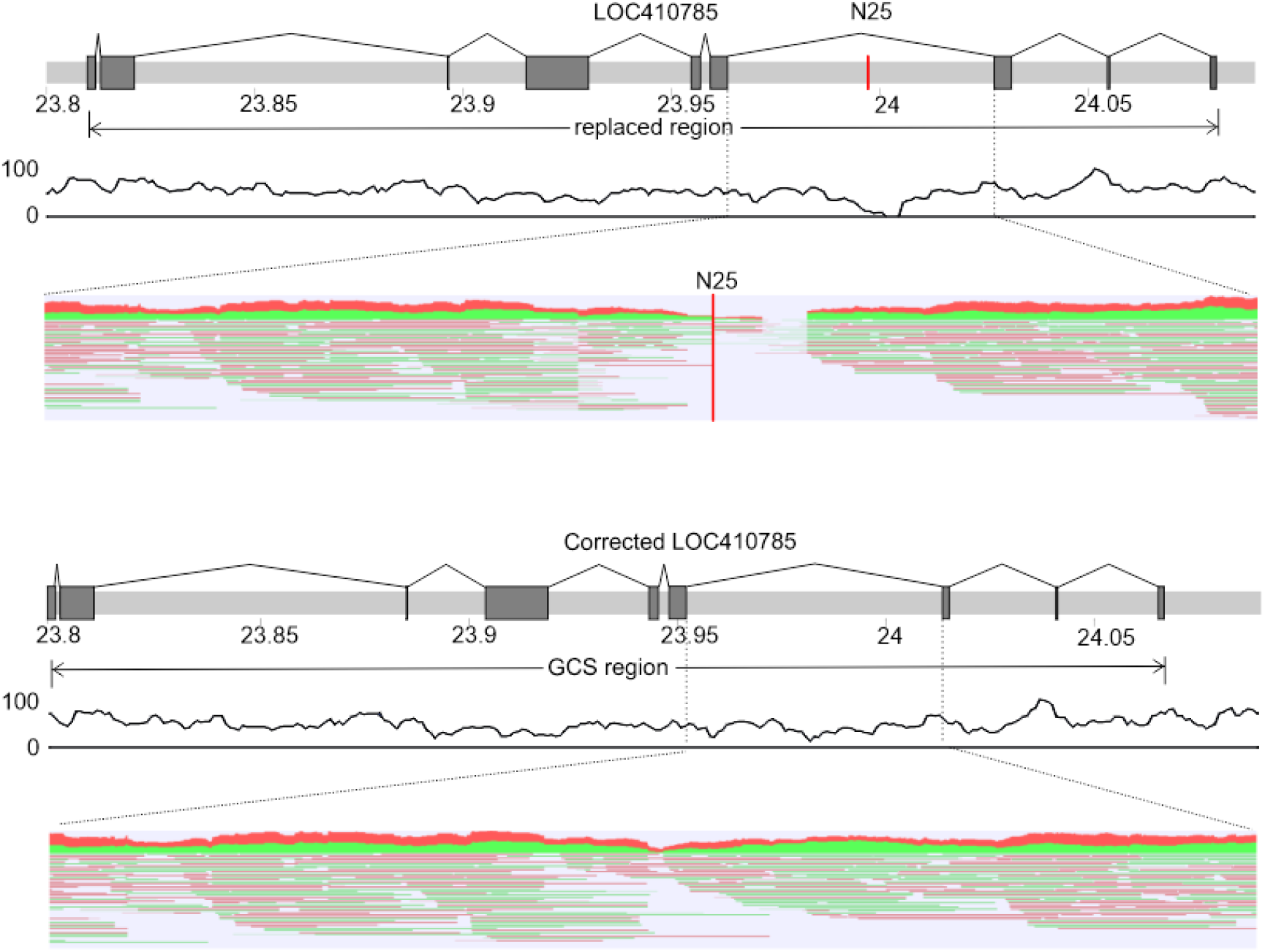
Gap-closing sequence from re-assembled Amel_HAv3_1 for gap 9 (N25) of chromosome 1. Exons are marked in dark gray. The red line N25 represents the gap. The black curve under the chromosomes shows PacBio reads coverage. Red-green hatching shows alignments of long PacBio reads to the zoomed region.

We failed to close some of the gaps using re-assembled Amel_HAv3_1 contigs alone. In such cases, we used sequences derived from the alternative assemblies INRA_AMelMel_1.0., ASM1384120v1 and ASM1384124v1. This allowed us to close two additional gaps. One of them is gap 2 of chromosome 1, which is located between LOC409701 and LOC113218996. For this gap, the GCSs were found in three alternative assemblies INRA_AMelMel_1.0., ASM1384120v1 and ASM1384124v1. These GCSs were aligned using the Kalign tool implemented in the Unipro UGENE (Okonechnikov *et al.* 2012). It should be noted that the GCSs from the ASM1384120v1 and ASM1384124v1 had the same repeat patterns, but minor sequence differences (UGENE Dotplot). Therefore, we selected GCSs from INRA_AMelMel_1.0 and ASM1384124v1 to create two corrected versions (Figure 4). To select one of them, we mapped PacBio reads using Minimap2 and found that the coverage in the ASM1384124v1 GCS was higher. We used this higher coverage version to close the gap. We then applied this approach to select the GCS for gap 1 in chromosome 3 (GCS source is ASM1384120v1). Details on genome coverage are given in Table S5.

**Figure 4.**
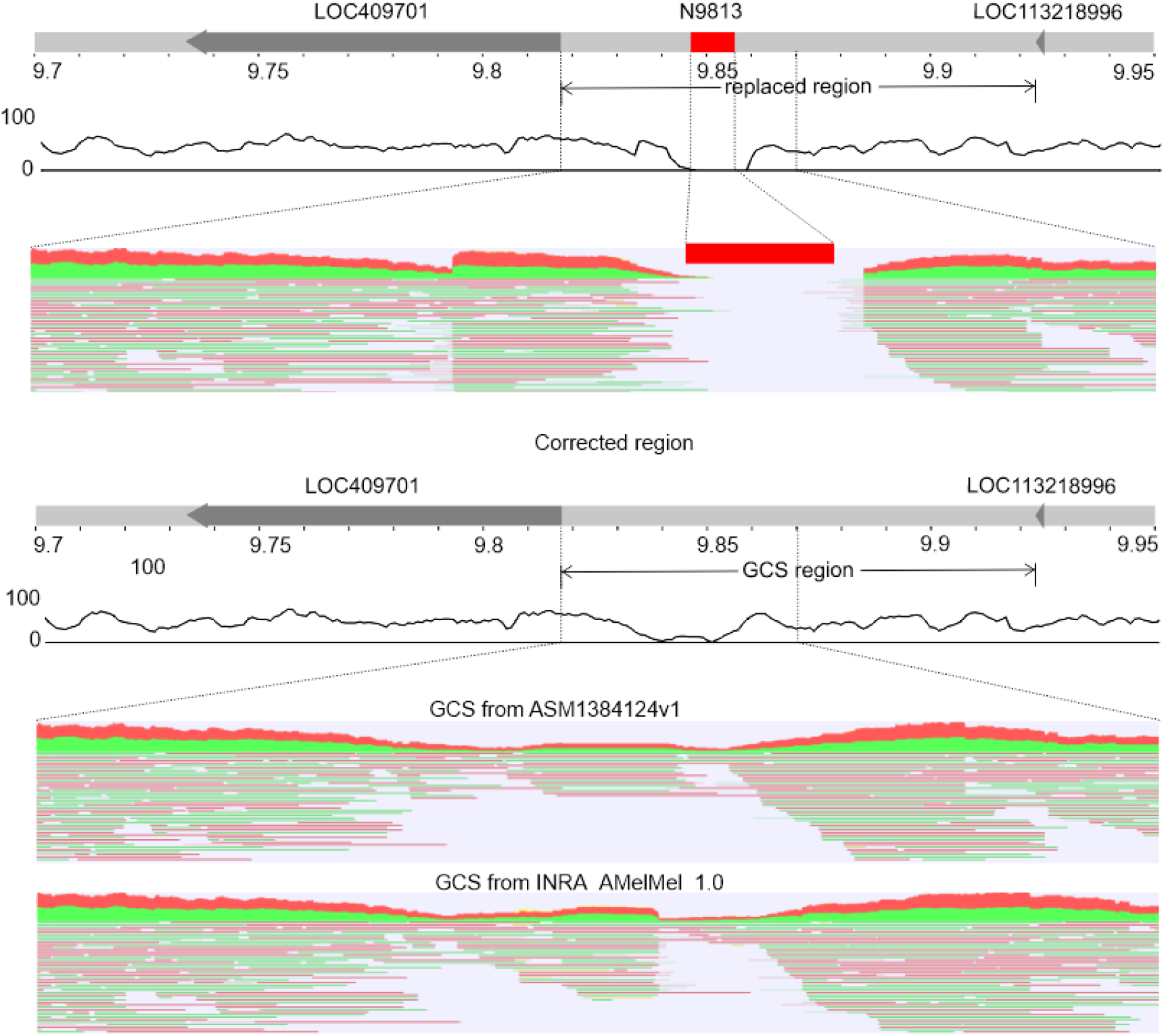
Gap closing sequence from ASM1384124v1 for the gap 2 of chromosome 1. The red square represents a gap, arrows - genes. The black curve under the chromosomes shows PacBio reads coverage. Red-green hatching shows alignments of long PacBio reads to the zoomed region.

### Positioning unplaced scaffolds

There are 11 chromosomes in Amel_HAv_3.1 that have unlocalized scaffolds and 43 unplaced/unlocalized scaffolds have genes. We determined the coordinates of these genes in the alternative assemblies. If the gene location, ordering, and orientation matched in more than two assemblies, we considered it to be the true location in the genome. Using this approach, we localized four unplaced scaffolds of the reference genome: NW_020555794.1 (40,528 bp, associated with chromosome 8, Figure 5), NW_020555815.1, and NW_020555816.1 (67,913 and 40,431 bp respectively, both associated with chromosome 10, Figure S2.9. and 3), and NW_020555860 (311,923 bp, Figure S6). Notably, two of these unlocalized scaffolds overlapped the gaps. The NW_020555794.1 closed the gap 1 in chromosome 8, and the NW_020555815.1 closed the gap 6 in chromosome 10 (Figure 5). The unplaced scaffold NW_020555860 along with the GCS from the corresponding alternative assembly was used to recover the proximal end of chromosome 16. We then mapped unlocalized scaffolds to the corrected reference using Minimap2 to validate their positioning.

**Figure 5.**
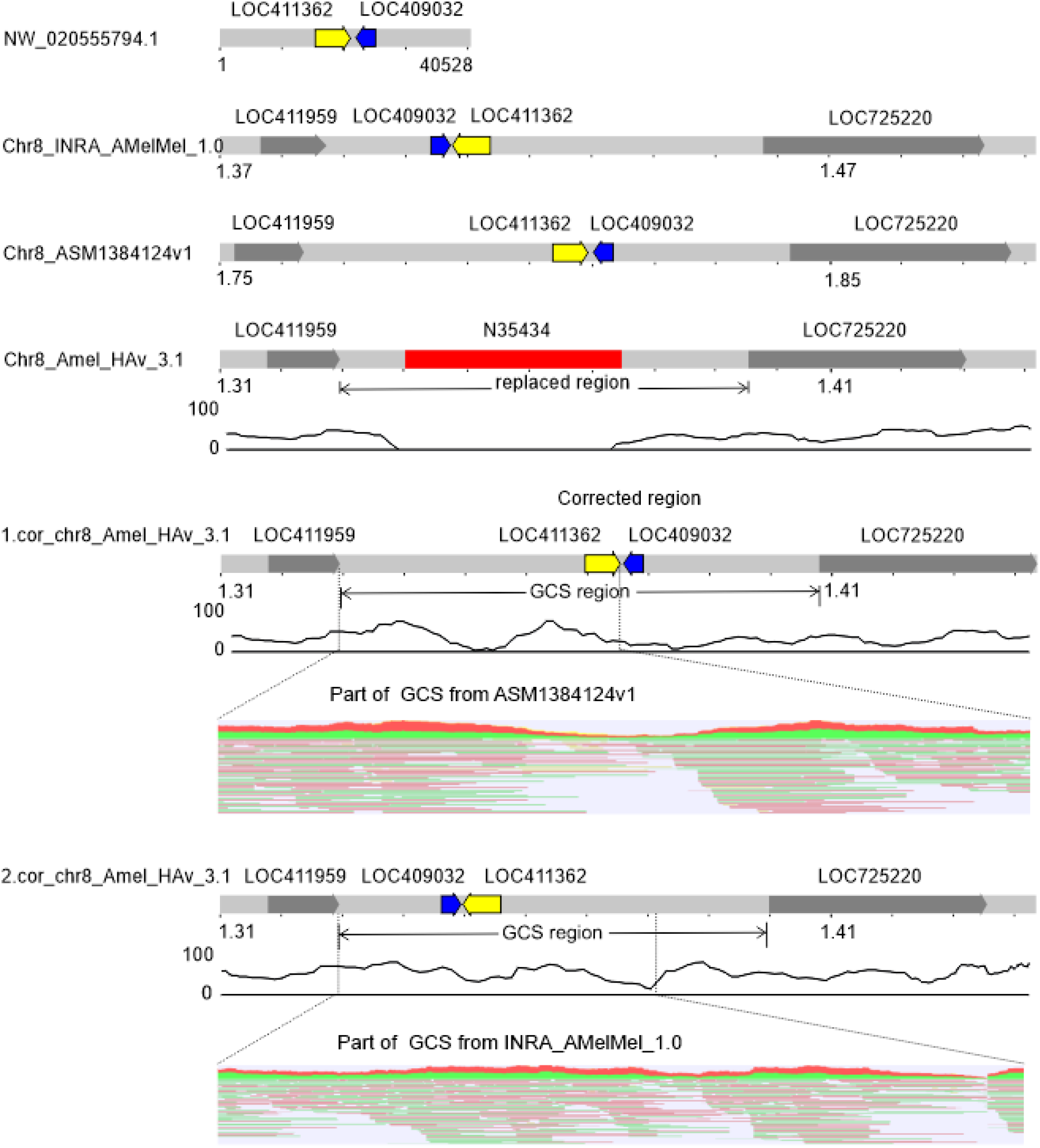
Gap-closing sequence from NW_020555794.1 for the gap 1 of chromosome 8. The red square represents gaps, and the arrows represent genes. The black curve under the chromosomes shows PacBio reads coverage. Red-green hatching shows alignments of long PacBio reads to the zoomed region. 1.cor_chr8_Amel_HAv_3.1 is a gap-closing sequence from ASM1384124v1, 2.cor_chr8_Amel_HAv_3.1 - from INRA_AMelMel_1.0.

Table 1 provides details of closed gaps and the corresponding GCSs. Six of the 13 gaps are located in genes and most of them have been closed by re-assembled Amel_HAv_3.1.

**Table 1.** Characteristics of gaps and corresponding GCSs.

| Gap (size, bp) | Replaced region (size, bp) | GCS source (size, bp) |
| --- | --- | --- |
| Chr1_gap2 (9,813) | from the end of LOC409701 to the start of LOC113218996 (106,010) | ASM1384124v1 (105,996) |
| Chr1_gap4 (1,978) | from the end of LOC414039 to start of LOC725387 (33,977) | Amel_HAv3_1_reFlye (34,179) |
| Chr1_gap6 (8,670) | LOC410685 (64,235) | Amel_HAv3_1_reND (52,788) |
| Chr1_gap8 (4,869) | LOC410674 (142,134) | Amel_HAv3_1_reFlye (142,302) |
| Chr1_gap9 (25) | LOC410785 (268,848) | Amel_HAv3_1_reFlye (266,084) |
| Chr2_gap1 (19,249) | from the end LOC102656216 to the start of LOC100577827 (128,592) | Amel_HAv3_1_reND (121,580) |
| Chr3_gap1 (25,238) | LOC410967 (145,799) | ASM1384120v1 (139,551) |
| Chr8_gap1 (35,434) | from end of LOC411959 to the start of LOC725220 (67,050) | ASM1384124v1 (78,460) |
| Chr8_gap3 (4,493) | from the start of LOC100578698 to the end of LOC100578828 (87,698) | Amel_HAv3_1_reFlye (92,821) |
| Chr8_gap4 (2,636) | AChE-2 (134,893) | Amel_HAv3_1_reND (134,907) |
| Chr10_gap6 (158,704) | from the start of LOC102654940 to the start of LOC409869 (200,539) | ASM1384124v1 (203,381) |
| Chr16_gap1 (56,203) | from the start of Mir993 to the start of LOC410648 (136,683) | Amel_HAv3_1_reND (142,661) |
| Chr16_gap2 (25) | LOC410655 (214,928) | Amel_HAv3_1_reFlye (221,629) |

### Telomere recovering and validation

The Amel_HAv_3.1 contains almost all distal telomeres, except the telomeres of chromosomes 5 and 11 (Figure 6). In chromosome 5 of the Amel_HAv_3.1, the distance between the last gene (LOC409500) and the end of the chromosome is 7,405 bp, while it is 19,481 bp in the INRA_AMelMel_1.0. Likewise, in chromosome 11 of the Amel_HAv_3.1, the distance between the LOC551454 and the end of the chromosome is 5,871 bp, while it is 21,258 bp in the INRA_AMelMel_1.0. Besides, INRA_AMelMel_1.0. has another gene (LOC113219342) that comes after LOC551454. In the Amel_HAv_3.1, the LOC113219342 is duplicated (Figure S4) and found in NW_020555814.1 (associated with chromosome 10) and NW_020555824.1 (13,259 bp, associated with chromosome 11). We used the telomeres of the alternative INRA_AMelMel_1.0 assembly to recover the telomeres lacking in the Amel_HAv_3.1 as shown in Figure 6. Then we mapped the PacBio reads to the corrected Amel_HAv_3.1 (Table 2).

**Figure 6.**
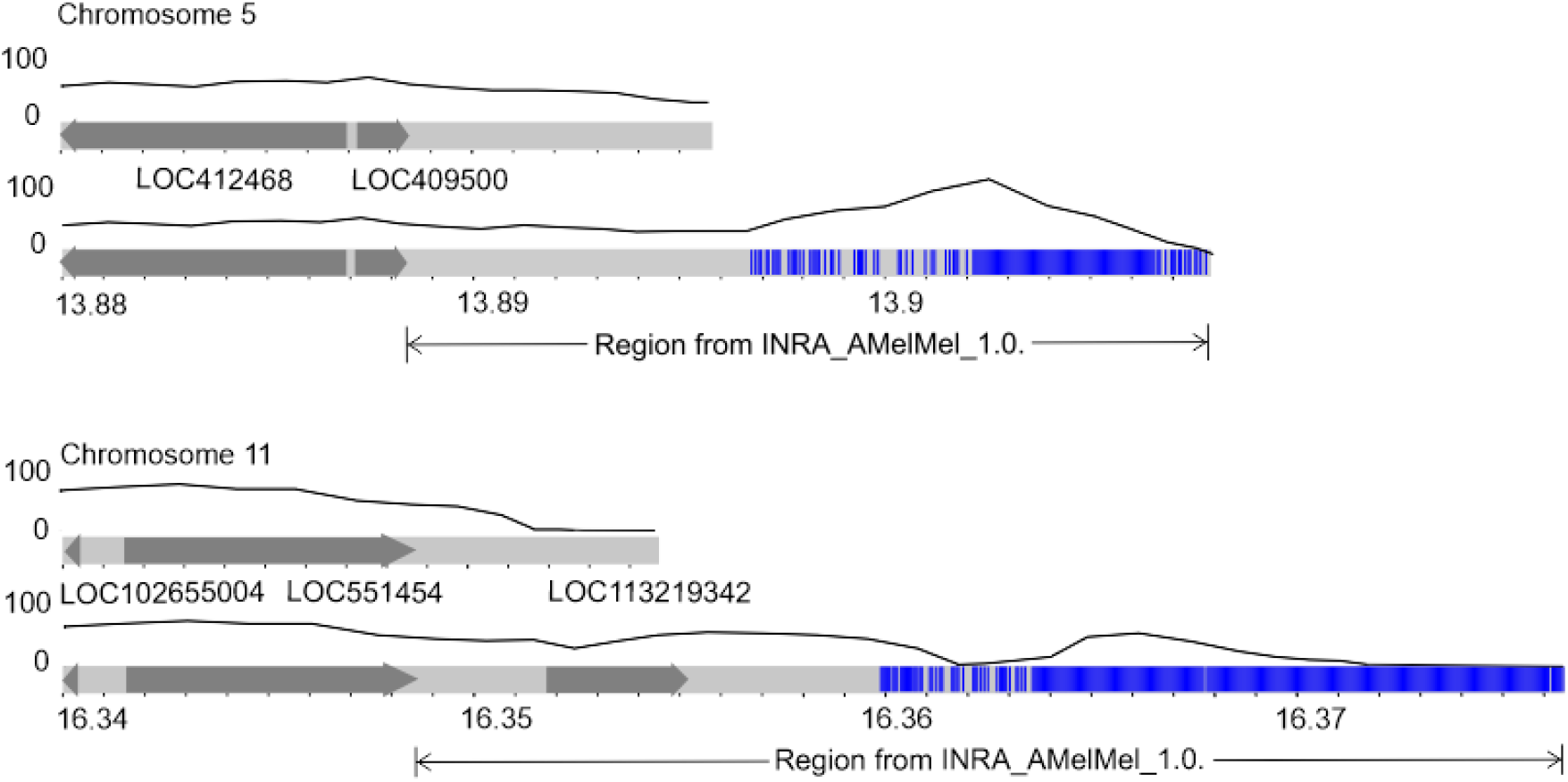
Distal ends of chromosomes 5 and 11 in the reference Amel_HAv_3.1 before (upper) and after (lower) correction with the mapped telomeric TTAGG motifs (blue), genes (dark gray), and PacBio reads coverage (black curve).

**Table 2.** Sequence coverage of the reference and corrected assemblies.

| Chr | Amel_HAv_3.1 |  |  | Corrected Amel_HAv_3.1 |  |  |
| --- | --- | --- | --- | --- | --- | --- |
|  | ID | Total read count | Average coverage | ID | Total read count | Average coverage |
| 1 | NC_037638.1 | 167,529 | 38.39 | cor_NC_037638.1 | 167,539 | 38.45 |
| 2 | NC_037639.1 | 96,598 | 38.16 | cor_NC_037639.1 | 96,696 | 38.23 |
| 3 | NC_037640.1 | 84,888 | 39.22 | cor_NC_037640.1 | 85,080 | 39.36 |
| 8 | NC_037645.1 | 75,555 | 37.85 | cor_NC_037645.1 | 75,835 | 38.02 |
| 10 | NC_037647.1 | 71,650 | 35.97 | cor_NC_037647.1 | 72,605 | 36.49 |
| 16 | NC_037653.1 | 43,400 | 38.32 | cor_NC_037653.1 | 45,829 | 38.50 |
| 5 | NC_037642.1 | 83,532 | 38.15 | cor_NC_037642.1 | 83,637 | 38.15 |
| 11 | NC_037648.1 | 100,362 | 39.19 | cor_NC_037648.1 | 100,433 | 39.16 |

### Redundancy removal and final corrected assembly assessment

To identify redundant sequences, we aligned unplaced/unlocalized scaffolds to the corrected reference genome using Minimap2. We found that scaffolds NW_020555860, NW_020555794.1, NW_020555815.1, NW_020555816.1, and NW_020555824.1 aligned to the replaced regions. Therefore, these scaffolds were determined to be redundant and deleted from the corrected Amel_HAv_3.1.

We ran BUSCO 4.0 with hymenoptera_odb10 and Liftoff to assess gene content in the corrected assembly (Table S6). The complete single-copy BUSCOs genes showed 0.4% increase, indicating a more complete assembly. Liftoff mapped all the reference genes, except the following three: LOC100578243, LOC113218760, and LOC113219414. These genes were, however, found to be in the genome using Minimap2 and represented duplicate genes (Figure S5).

We compared chromosome length (Table S7) and sequence coverage (Table 2) before and after gap closing. We observe improved coverage in almost all chromosomes except for chromosomes where telomeres have been added. Lack of improvement in such cases can be explained by the increased length of the chromosomes per number of reads.

## CONCLUSIONS AND PERSPECTIVES

This study presents a gap-closing effort in the honey bee reference genome using the assembly-to-assembly approach (Zhao *et al.* 2020). We began by re-assembling the Amel_HAv_3.1 using two different assemblers. The obtained re-assembled genomes as well as three alternative assemblies allowed us to find gap closing sequences and significantly improve the honey bee reference genome. We confirmed the accuracy of the corrected assembly by means of gene annotation and through mapping long PacBio reads. This approach has been successfully used for the human genome (Shi *et al.* 2016; Zhao *et al.* 2020).

Altogether, we closed 13 genomic gaps (327,337 bp) out of 51 and recovered two distal telomeres (47,356 bp). Our work fixed five unplaced scaffolds (474,054 bp in total) and produced 3 gapless chromosomes in the corrected Amel_HAv_3.1 reference. Our comparative analysis of honey bee genome assemblies suggests that assemblies based on PacBio reads failed in the same highly repetitive extended regions, notably on chromosome 10. Further work based on ultra-long Nanopore reads would be needed to fully resolve these extended repetitive regions.

Improving the reference genome of an organism is an important starting point in translating genomic information into its function at molecular, cellular, and organismal levels. We believe that our work on producing a more complete and accurate corrected_Amel_HAv_3.1 reference will facilitate novel downstream inferences in the field of honey bee research, which start with technical steps such as reference-guided scaffolding, marker/sequence mapping, and alike.

## ACKNOWLEDGMENTS

This study was supported by the Russian Foundation for Basic Research (project 19-54-70002) to A.N., U.Y., in part by the Ministry of Science and Higher Education of the Russian Federation (project No. AAAA-A21-121011990120-7) to M.K., Estonian Research Council grant PUT (PRG243), European Regional Development Fund (Project No. 2014-2020.4.01.16-0125), and ITMO University Fellowship to B.Y., and Eva Crane Trust Fund to B.H. and M.H.C.

## Author contributions

U.Y. conceived and designed the experiments. M.K. and U.Y. performed bioinformatics analyses. M.K. and R.A. designed artworks. M.K., B.Y., R.R., B.A.H., and U.Y. wrote the main manuscript text. A.N., B.A.H., M.H.C., and R.A. provided resources and laboratory space. All authors reviewed the manuscript.

All authors declare that they have no competing interests.

